# A panel of single nucleotide polymorphism (SNP) markers identifies potential duplicates in cassava (*Manihot esculenta* Crantz) varieties from Côte d’Ivoire

**DOI:** 10.1101/2021.05.24.445412

**Authors:** Edwige F. Yéo, William J-L. Amoakon, Justin S. Pita, J. Musembi Mutuku, Boni N’zué, Modeste K. Kouassi, Nasser Yao, Daniel H. Otron, Trushar Shah, Linda P. L. Vanié-Leabo, Kanh M. H. Kpahé, Raoul Sié, Fatogoma Sorho, Daouda Koné, Simon-Pierre A. N’guetta, Nazaire K. Kouassi, Morag E. Ferguson

**Author notes:** Corresponding author: **E-mail:** (JSP).

## Abstract

Accurate identification of varieties is paramount to optimizing efficiencies in the management and conservation of genetic resources. A relatively inexpensive, rapid methodology is required to identify putative duplicates from any collection, when morphological traits give insufficient discrimination. Here we select a panel of 36 SNPs, visualized using the Kompetitive Allele-specific PCR (KASP) system. We used a panel of 95 cassava genotypes from Côte d’Ivoire to identify varieties that are not duplicates and few potential duplicates which could be put forward for further verification. The genetic variability and population structure of the germplasm is also described. 36 SNPs were polymorphic across the panel of 95 varieties with polymorphic information contents ranging from 0.23 to 0.37. Using these SNPs, we were able to identify 66 unique genotypes from the panel of 95 genotypes, discriminate three sets of known duplicates and identify 11 sets of unknown putative duplicates which can be subjected to further verification using higher density genotyping. As expected in an outcrossing species, both expected heterozygosity (0.46) and observed heterozygosity (0.48) were high with an analysis of molecular variance (AMOVA) indicating that the majority of variation was within individuals. Three statistical approaches i.e., hierarchical ascending clustering, Bayesian analysis and discriminant analysis of principal components were used and all revealed low genetic differentiation between sub-populations, a conclusion that was supported by the low value of the fixation index (0.05). This panel of SNPs can be used to enhance cost-effectiveness and efficiency of germplasm conservation and enhance quality control at various stages in the breeding process through varietal tracking.

## Introduction

Breeding of improved varieties that meet specific product profiles for various uses and provide adaptation to different agro-ecologies and biotic stresses depend on the availability of well-curated and characterized genetic resources. This is particularly important in coping with the challenges posed by climate change [1]. A genotype of apparently little agronomic value today may become essential under the scenario of changing climate, diversified uses coupled with the appearance of new diseases [2]. It is paramount that crop genetic diversity is conserved and utilized as a key driver for securing further genetic improvement for sustainable development in the context of changing climate and population expansion [3].

Cassava is a staple food for millions of people around the world [4], yet the current diversity in the cultivated species *Manihot esculenta* Crantz is threatened by the replacement of a large number of genetically diverse landraces with a few improved varieties and the lack of adequate representation in international genebanks [5]. Other factors, such as disease pressure contribute to the loss of diversity. Apart from being predominantly clonally propagated, cassava is highly outcrossing, with random mating of gametes from distinct individuals at each generation, which generates substantial variation within individuals [6].

Germplasm repositories not only conserve germplasm, but should make it easily accessible in a disease-free condition, to plant breeders and researchers for utilization. The maintenance of *in vitro* germplasm repositories, often used for clonally propagated species, are however expensive. It is crucial that only unique accessions are maintained, with as much associated data as possible, such as passport data, characterization and evaluation data and farmer-knowledge. It is often difficult to discern whether a farmer-variety is unique when collecting in the field as the same genotype may have several different names in a given area [7, 8]. Accurate identification of cultivars/varieties could reduce the number of mislabeled clones and the cost of conservation [9]. Additionally, proper identification of varieties in crops is important for the varietal registration process, breeders seed production and trade [10].

Environmental conditions and different stages of plant development influence morphological descriptors [11, 12]. In addition, these tend to be limited in number. In Côte d’Ivoire, previous studies on the diversity of cassava varieties have focused on the use of agro-morphological traits [13, 14]. The quantitative morphological descriptors used were effective for selection in breeding, but could not fully elucidate genetic variability [8, 15]. Molecular markers have a much finer discriminatory power due to their relative abundance and the fact that they are not influenced by the environment. They enable the classification of genetic material using estimates of genetic distance and can also be used to quantify the relative proportion of ancestries derived from various founder genotypes of currently grown cultivars [16]. Among the molecular markers used for genotyping, single nucleotide polymorphisms (SNPs) have the advantage of being relatively low cost per generated data point. Their high abundance in the genome and their codominant state currently make them the most preferred marker [17, 18]. In cassava, SNP markers have been used to identify duplicate accessions in genebanks [6] and from field collections [19], in improved variety adoption studies [20] and assessing diversity [5, 21].

In this work, we identified (i) a low-density panel of SNPs suitable for varietal discrimination and fingerprinting in West African cassava germplasm (ii) unique varieties and putative duplicates in 95 cassava accessions from Côte d’Ivoire Cassava Germplasm Bank and cassava accessions from farmers’ fields and (iii) analyze the diversity and population structure of cassava varieties in this population using SNP markers. The common parameters of genetic diversity and genetic distance between pairwise accessions will allow respectively to explore the variability within the 95 accessions, the identification of unique varieties and putative duplicates using three different approaches to determine the genetic structure of these accessions i.e., Ascending Hierarchical Clustering (AHC), Discriminant Analysis of Principal Components (DAPC) and Bayesian analysis. The inference of the groups by AHC is based on the genetic distance between accessions while the other two methods infer the groups based on the membership coefficients in relation to common ancestors.

## Materials and Method

### Origin of the plant material

Ninety-five (95) accessions were used in this study and included 72 improved cassava varieties and 23 cassava landraces collected from farmers’ fields and germplasm from the Centre National de Recherche Agronomique (CNRA, Côte d’Ivoire), the International Institute of Tropical Agriculture (IITA, Nigeria) and Ghana (S1 table). The 72 improved cassava varieties included three different sets of known duplicates varieties (Bocou1(CM52)A, -B; TMS2 CNRA, -CSRS; Bocou2(188/00158)CNRA, -CSRS). The panel of the 95 varieties are currently conserved in open fields at the CNRA research station in Bouaké and the Centre Suisse de Recherche Scientifique (CSRS) research station in Bringakro, both located in central Côte d’Ivoire.

### Selection of SNPs

A sub-set of 36 SNP markers were selected from Expressed Sequence Tag (EST) derived SNPs by Ferguson [5, 22], and converted to KASP primers (LGC Biosearch technologies, UK) as a costeffective method for use in varietal identification and quality control. SNP markers were selected based on position (one from each arm of each of the 18 chromosomes) and Polymorphic Information Content (PIC) value above 0.365 within East African cassava germplasm [22].

### Genotyping

Sampling and sample shipment were done as per the LGC protocol. Leaf material was sampled from each cassava accession using the BioArk Leaf (from LGC Biosearch technologies) sample collection kit. The plate was sealed with a perforated (gas-permeable) heat seal and placed in a heavy-duty, sealed plastic bag with desiccant to dehydrate and preserve the leaf tissue during transit to LGC Biosearch technologies in the UK for DNA extraction and genotyping. Total genomic DNA was isolated from plant tissue using LGC’s Sbeadex™ DNA extraction, performed at LGC Biosearch technologies. Sbeadex is a magnetic bead-based extraction chemistry which uses a novel surface modification and two-step binding mechanism to allow tight binding of DNA, and a final pure water wash to give a high level of quality and purity. The 36 SNP markers genotyping was performed using the Kompetitive Allele-specific PCR system (KASP™) genotyping assays. KASP genotyping assays are based on competitive allele-specific PCR and enable bi-allelic scoring of SNPs and Insertions/ deletions at specific loci. The KASP genotyping assay consists of three components namely the sample DNA, KASP Assay mix and KASP Master mix. The SNP-specific KASP Assay mix and the universal KASP Master mix are added to DNA samples, a thermal cycling reaction is then performed, followed by an endpoint fluorescent read. The raw data analyzed and scored on a Cartesian plot, also known as a cluster plot in order to interpret the raw data and assigned a genotype to each DNA sample using LGC’s proprietary Kraken software. Results of genotyping were presented as homozygotes (A:A, C:C, G:G and T:T) and heterozygotes (A:T, A:C, A:G, C:A, C:T, C:G and G:T). Accessions and SNP markers with > 6% missing data were removed prior to diversity assessment. In addition, only one of each duplicate accession was retained.

### Analysis of genetic diversityEstimation of common genetic parameters

Polymorphic information content (PIC) is the potential of a marker to detect a polymorphism within a population [23]. A locus is considered polymorphic when the most frequent allele has a frequency of ≤ 0.95 [24]. PIC allows the determination of the informative capacity of a marker in a population from the allelic frequencies [25]. Its formula is 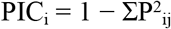, where *Pij* is the estimate of the frequency of genotype *j* at *i*th locus. Botstein classified PIC values as highly informative (PIC > 0.5), moderately informative (0.25 < PIC < 0.5) and less informative (PIC < 0.25) [26].

Expected heterozygosity (*He*) represents the theoretical rate of heterozygosity assuming the population meets the Hardy-Weinberg equilibrium (HWE). The *He* is calculated from the allelic frequencies according to the formula 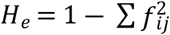; where *f_ij_* is the frequency of the *j*th allele of the *i*th locus. Observed heterozygosity (*Ho*) is the number of heterozygous individuals in relation to the total of individuals in the sample. It is calculated directly by the genotypic frequencies from the sample at a given locus *K*, according to the formula 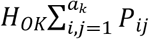 where *Pij* is the estimate of the frequency of genotype *ij* at locus *K* and *a_k_* is the number of alleles at locus *K*. *He* and *Ho* are ranged from 0 to 1 with 0 for no heterozygosity and 1 when there are many alleles at equal frequencies.

A genotype accumulation curve based on multi loci genotypes (MLGs) was used to determine the minimum number of SNPs needed to differentiate all unique Multi Loci Genotypes (MLGs).

### Genetic differentiation parameters (F-statistics)

The fixation index *Fit* is a measure of homozygosity of individuals in the total population. The fixation index, *Fis* [27] shows the differentiation of individuals within sub-populations (groups). It is calculated according to the formula: 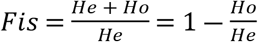. The fixation index *Fst* measures identity of individuals within sub-populations compared to individuals from other sub-populations within the total population. *Fst* = 1 – (*Hs/Ht*), where *Hs* is average of intra-population genetic diversity and *Ht* is genetic diversity across populations considered as a single population (total diversity). According to Wright: 0 < *Fst* < 0.05 is weak differentiation; 0.05 < *Fst* < 0.15 is moderate differentiation; 0.15 < *Fst* < 0.25 is significant differentiation; and *Fst* > 0.25 is very important differentiation [28]. These three parameters are linked as per the formula (1 – *Fit*) = (1 – *Fis*)(1 – *Fst*).

All parameters of genetic diversity and F-statistics were calculated with the HierFstat package 0.04-22 version [29] implemented in R version 3.3.3, with the exception of PIC which was calculated using PICcalc [30]. The HWE for each locus was calculated using the Adegenet package [31] implemented in R version 3.3.3. The genotype accumulation curve based on multi locus genotypes (MLGs) was performed using the Poppr package [32] also implemented in R version 3.3.3.

### Analysis of genetic structure

#### Variety identification and hierarchical ascending clustering based on Ward’s distance

A Ward’s minimum variance hierarchical clustering dendrogram was built from genetic distance using *plot.phylog* algorithm in the package Ade4 [33] as implemented in R version 3.3.3. The critical distance threshold to declare whether two accessions (varieties) are identical or not was based on the genetic distance between two representatives of the same accessions (duplicated previously for genotyping). Any two accessions whose genetic distance was below 0.05 (dissimilarity coefficient, Ward’s distance) were considered to be the same genotype. Dendrogram truncation was set using the *best.cutree* algorithm in the JLutils package [34] as implemented in R version 3.3.3 to highlight the genetic groups.

#### Bayesian analysis

The software STRUCTURE 2.3.4 version [35] was used to analyze the population structure of the cassava accessions. We used the Bayesian Markov Chain Monte Carlo (MCMC) approach based on the ADMIXTURE ancestry model which infers the genetic structure of populations while verifying the correct assignment of accessions to their group according to a probability Q [35]. The correlated allele frequencies model was applied in this analysis. The Bayesian approach assumes that the loci are in linkage equilibrium and that the sub-populations meet HWE requirements. STRUCTURE assumes that there are unknown K clusters, each of which is characterized by a set of allele frequencies at each locus [36]. The number of clusters was inferred using 15 independent runs for each value of K with 50,000 lengths of burn-in period and 500,000 MCMC replications after burn-in with K varying from 1 to 20. The best value of K (ΔK) was determined according to Evanno [37] using STRUCTURE HARVESTER 0.6.7 [38]. We used the probability Q matrix from the analysis to assign each accession to different clusters (K) using a critical level of probability Q at 70% for each one.

#### Discriminant analysis of principal components (DAPC)

The DAPC was performed using the Adegenet package [31, 39] as implemented in R version 3.3.3. This new approach provides the assignment of individuals to groups, a visual assessment of between population differentiation and a contribution of individual alleles to population structure. This method which combines principal components analysis and discriminant analysis (DA) is more suitable for populations that violate HWE and linkage equilibrium assumptions [31] such as in cassava which is a clonally-propagated crop. Unlike the STRUCTURE software, Adegenet software uses the non-model-based multivariate approach (that does not rely on HWE or assumes the absence of linkage disequilibrium). The DAPC assigns each individual to its home group according to a membership coefficient. In our work, the database was first transformed into a *genind* object. The number of principal components (PCs) and discriminant function that explained 98% of the total genetic variation were retained. To identify the optimal number of clusters (K), the *find-cluster* algorithm was used; this algorithm runs successive K-means clustering with increasing values of K. The lowest associated Bayesian Information Criterion (BIC) indicates the best number of clusters. The crossvalidation function *xvalDapc* was used to determine the correct number of PCs to be used and the number of discriminant functions to be saved to run the DAPC. The *xvalDapc* divides the data into two sets: training and validation sets with 90% and 10% of the data, respectively. The accessions of each group are selected by stratified random sampling, which ensures that at least one accession from each group in the original data is represented in both training and validation sets. The optimum number of PCs that should be retained is associated with the lowest root mean square error. Then the *dapc* algorithm was used to assign accessions into sub-populations. Contributions of the alleles to each discriminant function were highlighted by the loading plots.

#### Analysis of molecular variance (AMOVA)

AMOVA was performed to evaluate the distribution of genetic variation among the accessions using the package Poppr [32] implemented in R version 3.3.3. Before running AMOVA, the germplasm was grouped into different hierarchical levels, i.e., breeding patterns (improved varieties and landraces) and geographic origin of accessions (Nigeria, Côte d’Ivoire or Ghana) using the theoretical clusters obtained by the DAPC and Bayesian analysis.

#### Assessing the variability in relation to other cassava germplasm from Africa

To understand the extent of the variability of the accessions from Cote d’Ivoire in relation to other cassava germplasm from Africa, a selection of cassava from other African countries was added to the dataset and a combined analysis undertaken. The African cassava collection included 34 cassava accessions from Southern and Eastern Africa and two accession from West Africa which had previously been genotyped with the same 36 SNP markers (S2 Table). The combined and consolidated dataset of 111 cassava accessions were analyzed using Principal Coordinates Analysis (PCoA) and AHC, with the goal to estimate the extent of genetic similarity between Côte d’Ivoire germplasm and that from other African regions. To capture more variability, a 3-D PCoA was performed with the *cmdscale* function on the dissimilarity matrix constructed with the *vegdist* function of the vegan package [40] using the Bray-Curtis method. The plots were generated with the ggplot2 package [41] and plotly package [42].

## Results

### Seven SNPs can differentiate genotypes and identify putative duplicates

On the basis of the genetic distance threshold below 0.05, in the 95 accessions, we identified 66 unique genotypes, 10 pairs and one trio of unknown putative duplicate accessions and confirmed the three different sets of known duplicate varieties (Bocou1(CM52)A, -B; TMS2 CNRA, -CSRS; Bocou2(188/00158)CNRA, -CSRS). which were 25% of the total set of the 95 accessions. We found that two local varieties collected under the same local name were in fact different genotypes. For instance, a variety Yacé collected from CNRA and a variety collected in a farmer’s field under the same name were in fact different genotypes (Fig 1). Interestingly, a genotype accumulation curve based on MLGs found the minimum number of SNPs needed to differentiate all 66 unique MLGs is seven SNPs (S1 Fig).

**Fig 1.**
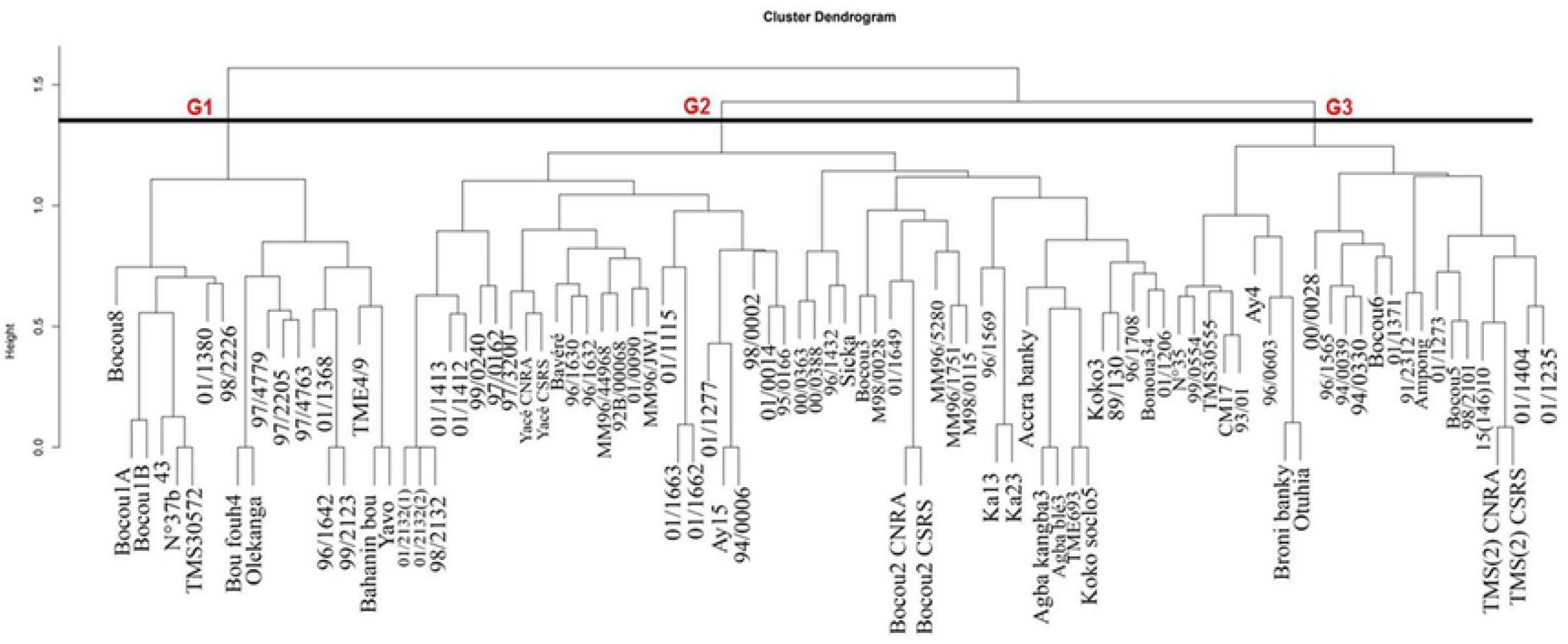
A dendrogram developed using Ward’s minimum variance method to show hierarchical clustering of the 95 cassava varieties reveals three groups (G1, G2 and G3).

### 36 SNP markers were polymorphic

The genomic positions and surrounding sequence of SNP markers used in this study are provided in Table 1. In the 95 accessions studied, all loci in the 36 SNP markers analyzed, were polymorphic. The marker Me.MEF.c.1094 had 16.7% of missing data so was removed from the panel of SNP markers for the genetic diversity assessment. The variety Bocou 8 had 44.4% of missing data, so was removed as well as 17 putative duplicates in the panel of accessions prior to genetic diversity analysis. Therefore, further analysis considered only 77 accessions (Table 2), which were used in the diversity and structure analysis comprising 66 unique genotypes and one of each of the putative duplicates. PIC values across the 77 unique accessions ranged from 0.23 to 0.37, with Me.MEF.c.1585 having 0.23, 17 markers having 0.37 and an average value of 0.35 (Table 2). All markers had PIC ? 0.30, excluding Me.MEF.c.1585 with PIC = 0.23. The *He* varied from 0.25 to 0.50, with Me.MEF.c.1585 having 0.25, 12 markers having 0.50 and an average of 0.46. In contrast, *Ho* ranged from 0.28 to 0.63, with Me.MEF.c.2268 having 0.28, Me.MEF.c.0284 and Me.MEF.c.1361 having 0.63 and an average of 0.48. The HWE analysis of six SNP markers showed that the rate of *Ho* was significantly different (*P* < 0.05) from that of *He*. For one of them i.e., Me.MEF.c.2268, this difference was highly significant (*P* < 0.001, Table 3).

**Table 1.**
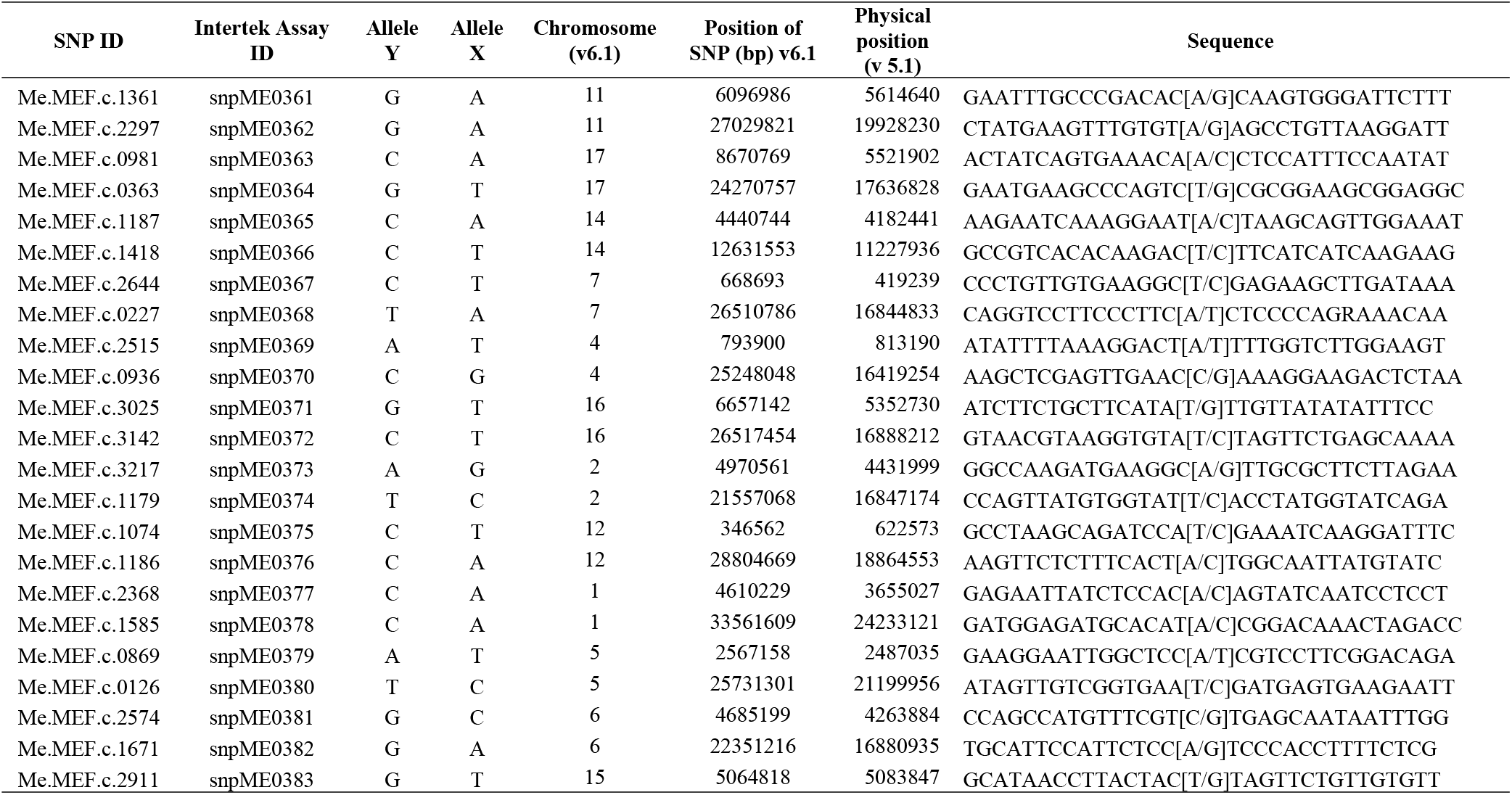
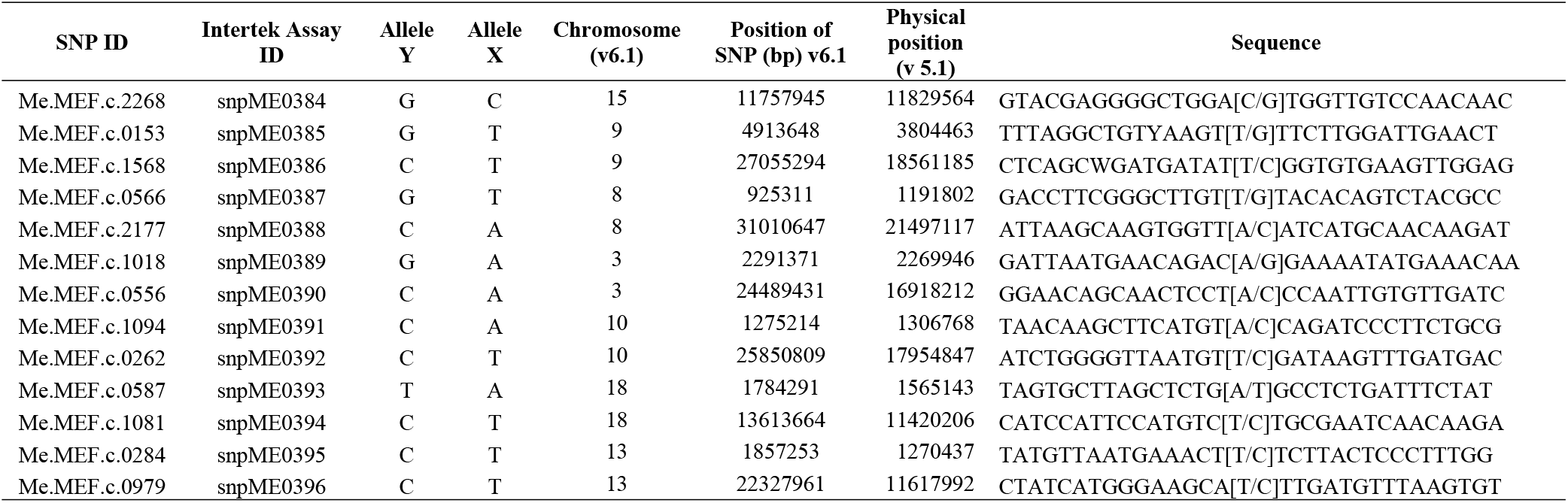
Genomic characteristics of 36 SNPs used in this study and their associated positions

**Table 2.**
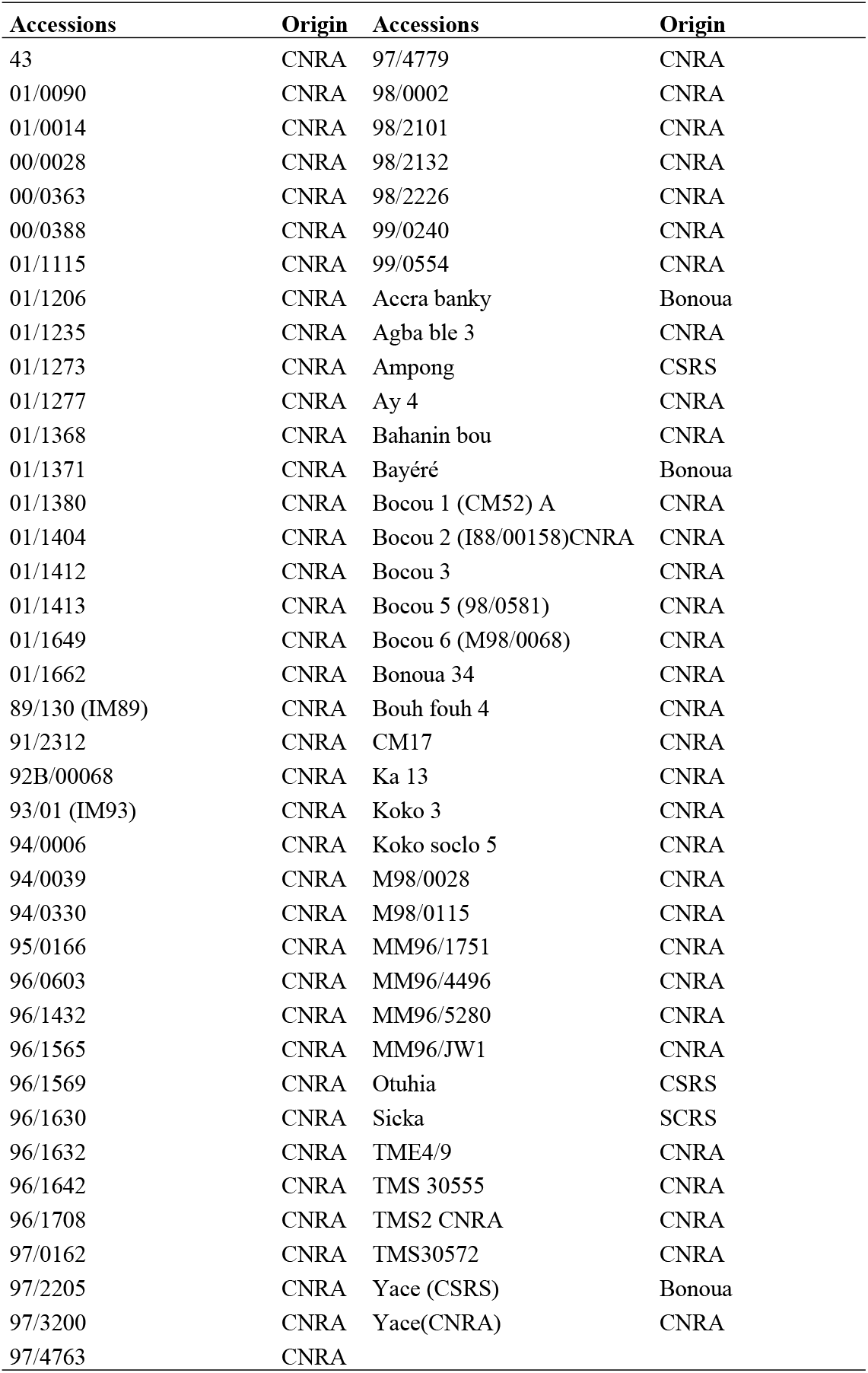
Origin of the 77 retained cassava varieties

**Table 3.**
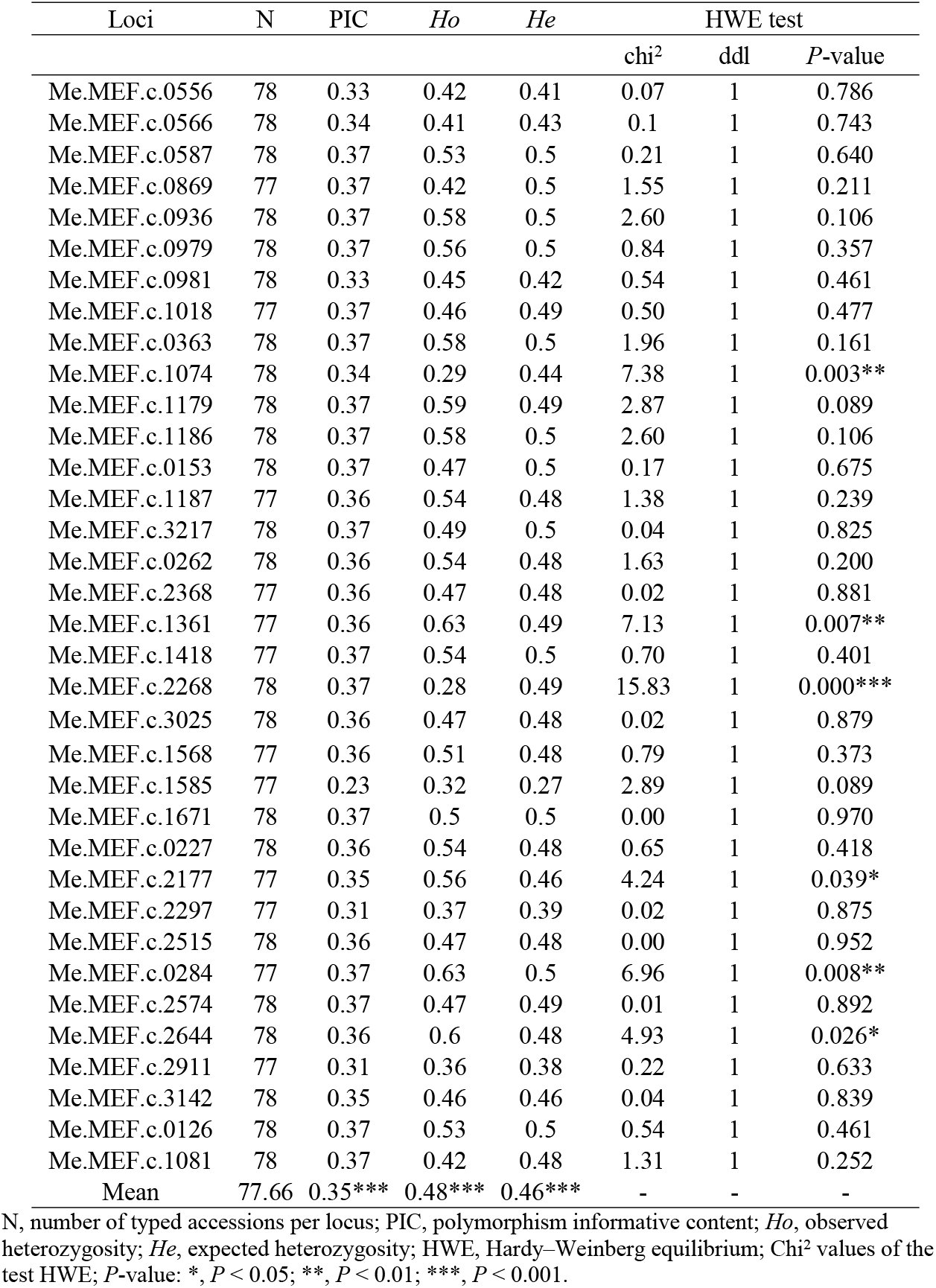
Genetic diversity parameters measured by locus from the Côte d’Ivoire germplasm

| Loci | N | PIC | *Ho* | *He* | HWE test |  |  |
| --- | --- | --- | --- | --- | --- | --- | --- |
|  |  |  |  |  | chi<sup>2</sup> | ddl | *P*-value |
| Me.MEF.c.0556 | 78 | 0.33 | 0.42 | 0.41 | 0.07 | 1 | 0.786 |
| Me.MEF.c.0566 | 78 | 0.34 | 0.41 | 0.43 | 0.1 | 1 | 0.743 |
| Me.MEF.c.0587 | 78 | 0.37 | 0.53 | 0.5 | 0.21 | 1 | 0.640 |
| Me.MEF.c.0869 | 77 | 0.37 | 0.42 | 0.5 | 1.55 | 1 | 0.211 |
| Me.MEF.c.0936 | 78 | 0.37 | 0.58 | 0.5 | 2.60 | 1 | 0.106 |
| Me.MEF.c.0979 | 78 | 0.37 | 0.56 | 0.5 | 0.84 | 1 | 0.357 |
| Me.MEF.c.0981 | 78 | 0.33 | 0.45 | 0.42 | 0.54 | 1 | 0.461 |
| Me.MEF.c.1018 | 77 | 0.37 | 0.46 | 0.49 | 0.50 | 1 | 0.477 |
| Me.MEF.c.0363 | 78 | 0.37 | 0.58 | 0.5 | 1.96 | 1 | 0.161 |
| Me.MEF.c.1074 | 78 | 0.34 | 0.29 | 0.44 | 7.38 | 1 | 0.003** |
| Me.MEF.c.1179 | 78 | 0.37 | 0.59 | 0.49 | 2.87 | 1 | 0.089 |
| Me.MEF.c.1186 | 78 | 0.37 | 0.58 | 0.5 | 2.60 | 1 | 0.106 |
| Me.MEF.c.0153 | 78 | 0.37 | 0.47 | 0.5 | 0.17 | 1 | 0.675 |
| Me.MEF.c.1187 | 77 | 0.36 | 0.54 | 0.48 | 1.38 | 1 | 0.239 |
| Me.MEF.c.3217 | 78 | 0.37 | 0.49 | 0.5 | 0.04 | 1 | 0.825 |
| Me.MEF.c.0262 | 78 | 0.36 | 0.54 | 0.48 | 1.63 | 1 | 0.200 |
| Me.MEF.c.2368 | 77 | 0.36 | 0.47 | 0.48 | 0.02 | 1 | 0.881 |
| Me.MEF.c.1361 | 77 | 0.36 | 0.63 | 0.49 | 7.13 | 1 | 0.007** |
| Me.MEF.c.1418 | 77 | 0.37 | 0.54 | 0.5 | 0.70 | 1 | 0.401 |
| Me.MEF.c.2268 | 78 | 0.37 | 0.28 | 0.49 | 15.83 | 1 | 0.000*** |
| Me.MEF.c.3025 | 78 | 0.36 | 0.47 | 0.48 | 0.02 | 1 | 0.879 |
| Me.MEF.c.1568 | 77 | 0.36 | 0.51 | 0.48 | 0.79 | 1 | 0.373 |
| Me.MEF.c.1585 | 77 | 0.23 | 0.32 | 0.27 | 2.89 | 1 | 0.089 |
| Me.MEF.c.1671 | 78 | 0.37 | 0.5 | 0.5 | 0.00 | 1 | 0.970 |
| Me.MEF.c.0227 | 78 | 0.36 | 0.54 | 0.48 | 0.65 | 1 | 0.418 |
| Me.MEF.c.2177 | 77 | 0.35 | 0.56 | 0.46 | 4.24 | 1 | 0.039* |
| Me.MEF.c.2297 | 77 | 0.31 | 0.37 | 0.39 | 0.02 | 1 | 0.875 |
| Me.MEF.c.2515 | 78 | 0.36 | 0.47 | 0.48 | 0.00 | 1 | 0.952 |
| Me.MEF.c.0284 | 77 | 0.37 | 0.63 | 0.5 | 6.96 | 1 | 0.008** |
| Me.MEF.c.2574 | 78 | 0.37 | 0.47 | 0.49 | 0.01 | 1 | 0.892 |
| Me.MEF.c.2644 | 78 | 0.36 | 0.6 | 0.48 | 4.93 | 1 | 0.026* |
| Me.MEF.c.2911 | 77 | 0.31 | 0.36 | 0.38 | 0.22 | 1 | 0.633 |
| Me.MEF.c.3142 | 78 | 0.35 | 0.46 | 0.46 | 0.04 | 1 | 0.839 |
| Me.MEF.c.0126 | 78 | 0.37 | 0.53 | 0.5 | 0.54 | 1 | 0.461 |
| Me.MEF.c.1081 | 78 | 0.37 | 0.42 | 0.48 | 1.31 | 1 | 0.252 |
| Mean | 77.66 | 0.35*** | 0.48*** | 0.46*** | - | - | - |

N, number of typed accessions per locus; PIC, polymorphism informative content; *Ho*, observed heterozygosity; *He*, expected heterozygosity; HWE, Hardy–Weinberg equilibrium; Chi<sup>2</sup> values of the test HWE; *P*-value: \*, *P* < 0.05; \*\*, *P* < 0.01; \*\*\*, *P* < 0.001.

### Genetic differentiation

Analysis of genetic differentiation was conducted on the 95 cassava accessions and was not significant (*Fit* = −0.03, *P* = 0.29). Of the 35 loci, 14 had a positive inbreeding coefficient (*Fis*) value. However, the *Fis* mean value across all 35 loci was highly significant (−0.09, *P* = 0.001). Negative fixation index (*Fis*) values were estimated for 25 loci, and positive values were observed for the 10 others. Overall, the genetic differentiation between groups (sub-populations), taking into account all 35 loci, was weak with *Fst* = 0.05 and highly significant (*P* < 0.001, S3 Table).

### Analysis of genetic structure: Duplicate accessions and missing data may bias population structuring

#### Analysis of the total set of the 95 cassava accessions

The number of groups within the total set of the 95 cassava accessions varied according to each of the three analysis approaches applied in this study. The dendrogram obtained from the hierarchical ascending clustering identified three groups when using a level of dissimilarity coefficient of about 1.4 according to the algorithm best.cutree (Fig 1). However the optimal number of groups, determined according to the software STRUCTURE was two (S2 Fig); this coincided with the highest value of ΔK from the Evanno method (S3 Fig). Finally, the total set of the 95 cassava accessions were clustered into five groups in accordance with the lowest BIC value from the DAPC analysis (S4 Fig). The scatter plot of the DAPC show the representation of the five groups (S5 Fig).

#### Analysis of the set of the retained 77 cassava accessions

The dendrogram resulting from the hierarchical ascending clustering showed three groups when using a level of dissimilarity coefficient of about 1.5 according to the algorithm *best.cutree*. This grouping corresponded to that from the STRUCTURE-like analysis using the ADMIXTURE program to assign individuals proportionally to hypothetical founder populations. Each of the main branches of the dendrogram formed a distinct ancestry group highlighted by the barplot from the STRUCTURE software (Fig 2) which represents the estimated ancestries (Q). Thus, the optimal number of groups, determined according to the ADMIXTURE program was three for the 77 retained cassava accessions in coincidence with the highest value of ΔK from the Evanno method (S6 Fig). This result was validated by the DAPC method that is considered free of Hardy-Weinberg and linkage disequilibrium assumptions. In accordance with the lowest BIC value from the DAPC analysis, the 77 retained cassava accessions were grouped in three groups (S7 Fig). A major difference between the results of the latter two clustering methods was the propensity of the DAPC analysis to assign entire individuals to a single cluster compared to ADMIXTURE program, which was able to assign admixed individuals to multiple clusters. Thus, the membership coefficient of the accessions varied from 29% to 74% for ADMIXTURE program while it varied from 80% to 100% for the DAPC analysis (S7 and S8 Figs). All the cassava genotypes had their ancestry traced back to at least one of the three sub-populations from ADMIXTURE program. The clusters from DAPC mostly corresponded to sets of genetically similar groups of admixed individuals that shared the same ancestries (S4 Table). The scatter plot of the DAPC shows the representation of the three groups from the retained 77 accessions (Fig 3). According to the loading plots (S9 Fig) the locus with the most contribution was Me.MEF.c.2574 (0.10) for axis 1. For axis 2, most of the contributions was Me.MEF.c.2268 (0.17). These alleles best describe the variability of the population and optimally discriminate the variability existing between sub-populations.

**Fig 2.**
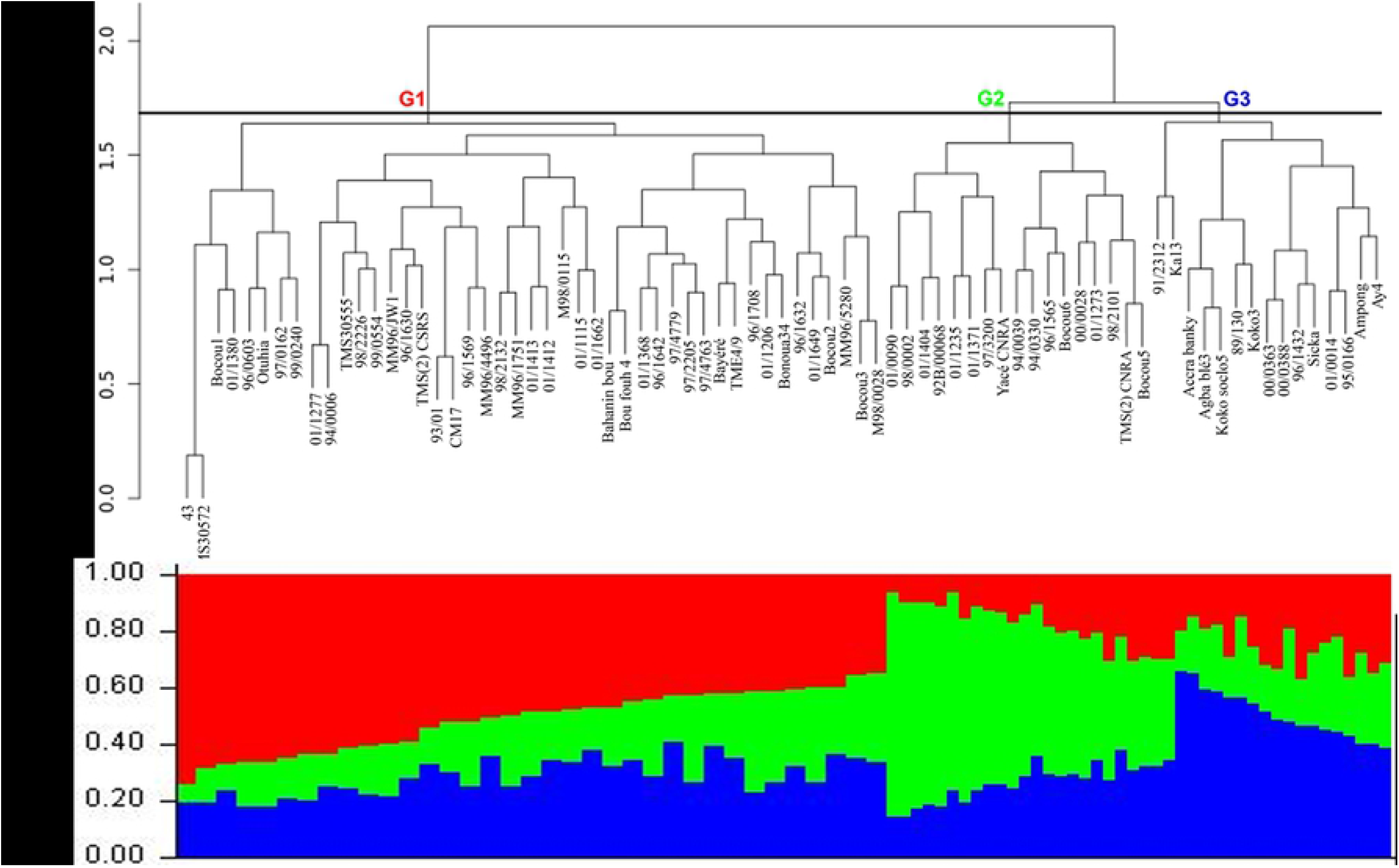
(A) A dendrogram developed using Ward’s minimum variance method to show hierarchical clustering of the 77 retained cassava varieties revealed three groups (G1, G2 and G3). (B) The main branches of the dendrogram correspond to a distinct ancestry group (Red, green and blue) highlighted by the barplot from the STRUCTURE software. Each accession is represented by a vertical bar. The membership coefficient of the accessions varied from 29% to 74%

**Fig 3.**
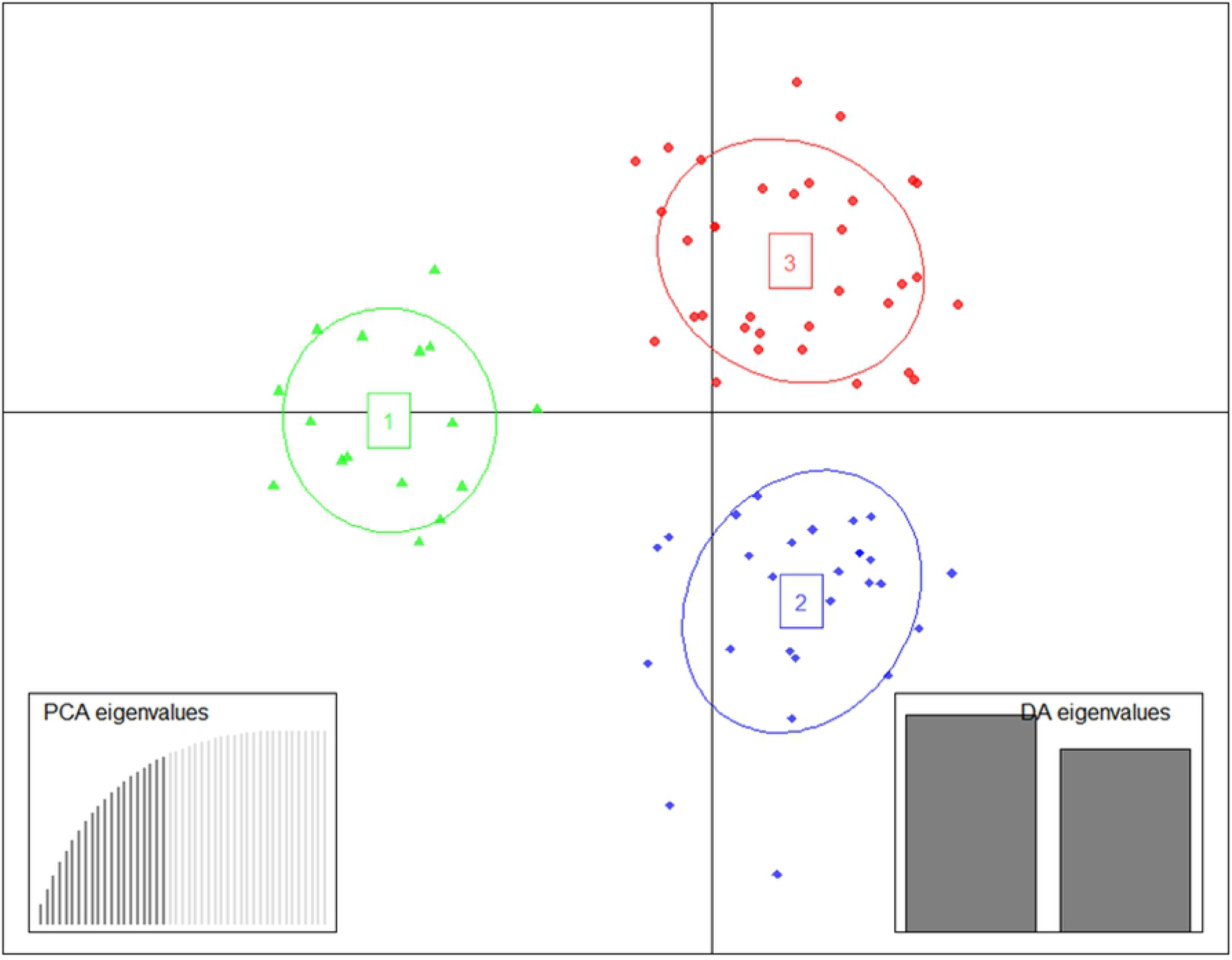
Plot of Discriminant analysis of principal components (DAPC) for three assigned genetic, clusters from the 77 retained varieties, each indicated by different colors. Dots represent different varieties. Inset left bottom corner and inset right bottom corner, show the eigenvalues of the 15 principal components and the eigenvalues of the two discriminant functions retained for the analysis respectively

### Maximum Molecular variance revealed by SNPs exists within individuals

AMOVA is reported against four levels of clustering formed based on *a priori* information (breeding patterns and geographical origin) and *a posteriori* information i.e., theoretical clusters obtained with DAPC and STRUCTURE. We found that the most significant differences in the molecular variance of the SNPs existed within individuals for all hierarchical levels ranging from 99.63% to 99.65% for STRUCTURE groups and geographical origins respectively with an intermediate value of 99.64% for breeding pattern and DAPC group. Likewise, the variation between (0.01–0.09%) and within populations (0.27–0.34%) was low for the four levels of clustering. The variation between populations varied between 0.01 and 0.09% for geographical origins and DAPC group respectively while the variation within population ranged from 0.27% for DAPC group to 0.34% for breeding pattern and geographical origins (Table 4).

**Table 4.** AMOVA considering two groups according to breeding patterns, four groups according to geographic origins, five groups according DAPC and two groups according to STRUCTURE

| Source of variation | Improved varieties and landraces |  |  | Geographical origins |  |  |
| --- | --- | --- | --- | --- | --- | --- |
|  | Df | Mean square | % of variation | Df | Mean square | % of variation |
| Between population | 1 | 37.20 | 0.02 | 4 | 29.07 | 0.01 |
| Within population | 93 | 15.59 | 0.34 | 90 | 15.23 | 0.34 |
| Within individuals | 95 | 17.54 | 99.64 | 95 | 17.56 | 99.65 |
| Source of variation | DAPC groups |  |  | STRUCTURE groups |  |  |
|  | Df | Mean square | % of variation | Df | Mean square | % of variation |
| Between population | 4 | 82.81 | 0.09 | 1 | 99.55 | 0.04 |
| Within population | 90 | 12.84 | 0.27 | 93 | 14.92 | 0.33 |
| Within individuals | 95 | 17.54 | 99.64 | 95 | 17.54 | 99.63 |

### Variability within the 95 accessions spans across the variability from other regions in Africa

PCoA was unable to distinguish clear groupings of the combined set of data of African accessions with those from Cote d’Ivoire using three dimensions accounting for 30.50% of the variation (Fig. 4). Likewise, the dendrogram showed three closely related groups with the 34 added cassava from others region in Africa being distributed throughout the three groups (Fig. 5).

**Fig 4.**
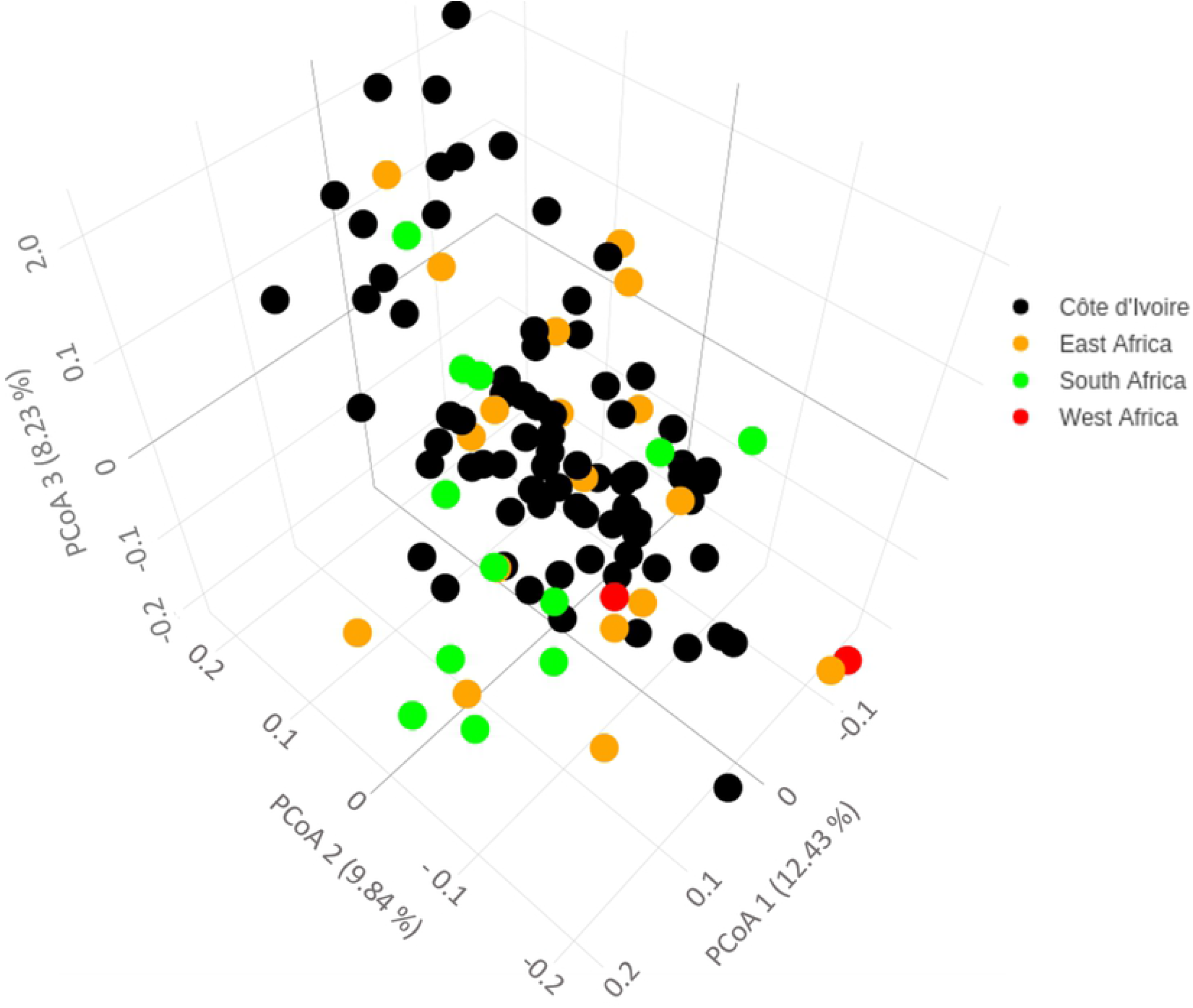
Principal Coordinates Analysis (PCoA) on the Côte d’Ivoire germplasm and other African germplasm genotyped using the 36 SNPs showing the level of relatedness and diversity among the populations. Dots represent different accessions. Black color represents the accessions from Côte d’Ivoire; Green, Orange and Red colors those from South, East, and West Africa respectively.

**Fig 5.**
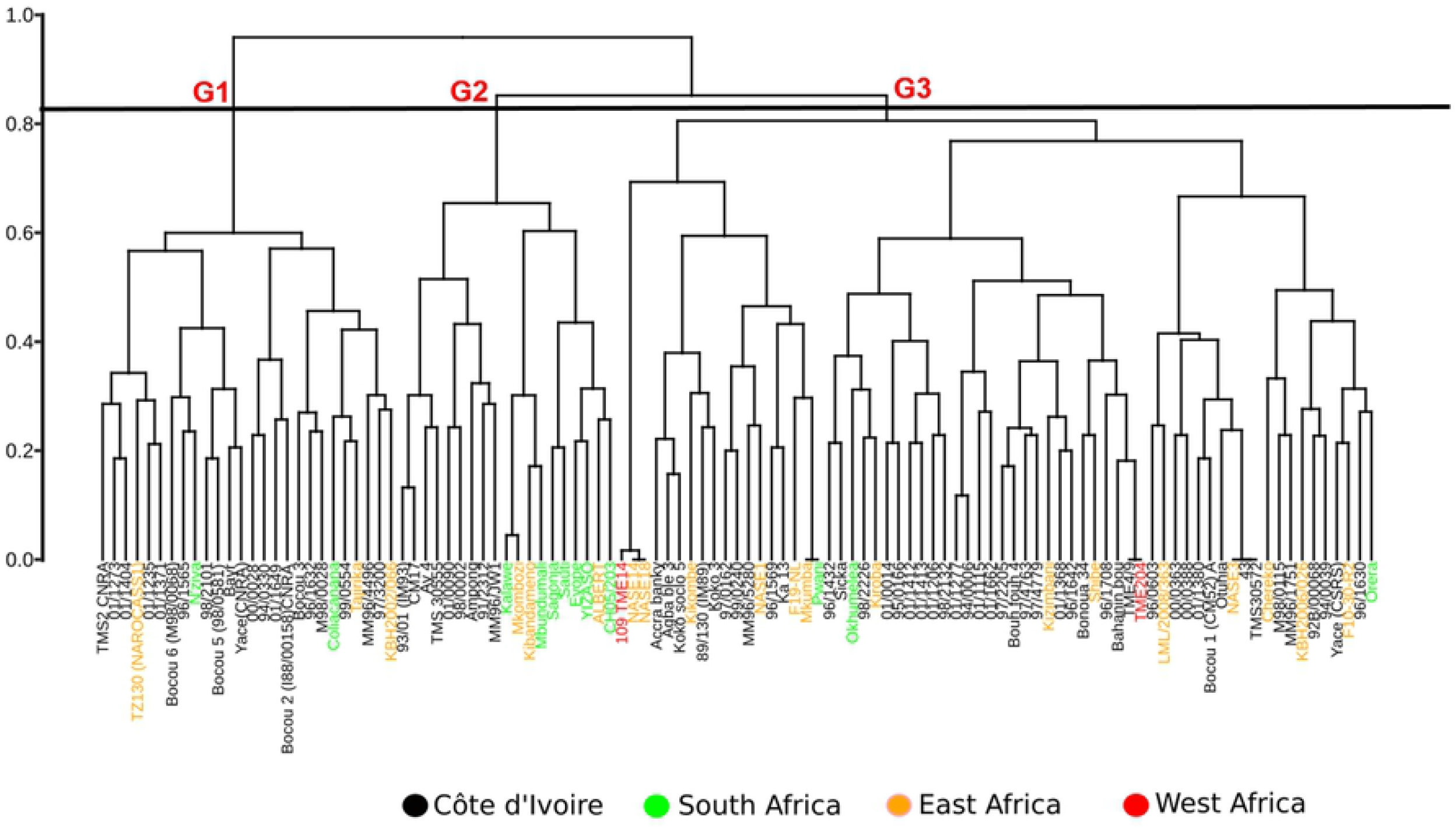
Dendrogram developed using Ward’s minimum variance method to show hierarchical clustering of the 111 combined cassava accessions revealing that. the 34 added cassava from others region of Africa are distributed throughout the three groups **(G1, G2 and G3)**. Black color represents the accessions from Côte d’Ivoire; Green, Orange and Red colors those from South, East, and West Africa respectively.

## Discussion

A polymorphism rate of 100% was obtained for the 95 accessions using the 36 SNP markers implemented in this study. These results confirm the effectiveness of these loci to fingerprint the studied accessions. With the exception of locus Me.MEF.c.0869 with PIC = 0.23, hence less informative and Me.MEF.c.1094 which was removed in the panel of SNP markers due to 16.7% of missing data, all other loci had PIC values of 0.31–0.37 and were highly informative. The SNPs were initially selected based on PIC values of a predominantly East African germplasm panel [5; 22]. Due to some population differentiation between West African and East African germplasm [5], it was important to validate these SNPs for further use in West African germplasm. In this study, we validated these SNPs. The SNPs are available either through LGC Biosearch technologies or the High-throughput Genotyping platform (HTPG) at the International Crops Research Institute for the Semi-Arid Tropics (ICRISAT). Interestingly, the average PIC value of 0.35 we observed in this study is higher than the 0.26 obtained by Ferguson [22]. This difference is likely due to the lower number of accessions investigated in the previous studies compared to the current study. It is notable that other types of markers such as simple sequence repeats (SSR) give much higher PIC values in cassava. For example, PIC values up to 0.75 with an average of 0.53 have been reported by Asare [43]. This difference is likely due to the bi-allelic nature of SNP markers compared to multi-allelic SSRs. The *He* and *Ho* values ranged from 0.25–0.50 and 0.29–0.64, respectively, with corresponding averages of 0.46 and 0.49 (Table 2). These values could indicate a high diversity within the 95 accessions analyzed. This was confirmed by the PCoA and AHC performed on a dataset combining Côte d’Ivoire germplasm and a collection of cassava germplasm from other African regions and clearly showed that the diversity within these accessions is similar to the variability at a continental level (Figs 4 and 5). These markers were unable to discriminate germplasm from West Africa from that of East Africa, as has been found in previous studies [5] based on 1,124 SNPs. It is likely that this is due to the limited number of SNPs used in this study.

Our findings revealed that six of the 35 loci analyzed significantly deviated from HWE (Table 2) and four were due to an excess of heterozygotes confirming the presence of a high genetic diversity that could be attributed partly to the presence of improved varieties which were obtained from the multiple crosses conducted by IITA and CNRA, and partly due to the natural hybridizations that occur in this strongly outcrossing species in farmers’ fields. In fact, plants from these natural hybridizations are often selected by farmers if they appear to be vigorous [44]. Through this action, they indirectly select genotypes which contribute to increased genetic variability in fields, as well as to diversity in the next generation of cassava seed in the field [12]. The *Fit* mean value of −0.03 across all loci indicated a non-significant deficit of homozygotes of 3% in the global population of accessions. The *Fis* mean value of −0.09 indicated a significantly higher excess of heterozygotes inside sub-populations when taken individually. Moreover, the relatively low value of *Fst* (0.05) indicates a low genetic differentiation between sub-populations (Table 3). Therefore, much of the genetic variability within the accessions is explained by the variation within individuals.

The synonymy revealed by the analysis of the accessions collected under different names but with the same genotype, could be explained by the plasticity of the morphological characters and/ or farmers giving new names as a genotype is introduced to a community as observed by Elias [45]. We identified 66 unique genotypes from the panel of 95 genotypes and 11 sets of unknown putative duplicates which we propose should be subjected to verification using higher density genotyping. These findings show the possible existence of the same cultivar under several entry numbers in the Côte d’Ivoire Cassava Gene Bank conserved at CNRA.

In this study, Ascending Hierarchical Clustering highlighted three genetic groups (Fig 1). From previous studies that used this method, we have learned that the cassava germplasm of Côte d’Ivoire can be structured into eight groups based on morphological characters [13, 46]. However, the absence of perfect congruence between morphological and molecular data revealed by Pissard [47] suggests that the morphological data can be useful for highlighting morphotypes but is not appropriate for studying genetic structure. The three methods of clustering used detected the same number of groups for the 77 retained cassava accessions, showing that the presence of the putative duplicate accessions and the missing data over 6% biased the genetic structuring for the total set of the 95 cassava accessions. The dendrogram allowed us to efficiently classify accessions according to the genetic distance between them and also to highlight the putative duplicate accessions. Knowledge of genetic proximity is important for genetic crosses in order to maximize efficient hybridization. However the ancestry information is important since it provides a framework for determining the contribution of specific germplasm in development of new varieties and therefore show indirect impact of germplasm originating from a specific breeding program [48]. This was achieved through the analysis of the populations structure from the ADMIXTURE program and DAPC analysis. Although we obtained the same number of groups with DAPC and ADMIXTURE program the latter method revealed large number of individuals with two or more ancestries while DAPC analysis mostly assigned individuals to single clusters. According to Jombart [31], the type of population structure influences the precision of the method. The inferences in structured populations in the discontinuous population structure such as island model are more precise than in continuous populations, which seems to be the case for the cassava germplasm which has complex population structure [15, 49]. The contribution of alleles to the groupings identified by DAPC allows the identification of genomic regions that drive genetic divergence among groups [29]. However, AMOVA analysis showed that the variation between and within populations was low (Table 4). These results show that the populations were not clearly structured, and consequently the sub-populations did not vary from each other. This could also be interpreted as suggesting that there was little variation in allele frequencies between groups. This limited differentiation among groups is likely due to the 1) limited number of bi-allelic SNPs used, 2) frequent movement of improved varieties between breeding centers such as IITA and CNRA and 3) farmers being conservative in using the same varieties over a long period of time. The latter reason is also reinforced by a poor variety replacement strategy by breeding institutions in Africa.

This study contributes to our current understanding of the merits of using molecular markers to analyze genetic structure. Indeed, Kawuki [15] showed that there is limited power of discrimination of cassava accessions based on morphological descriptors when evaluating the phenotypic variability of the cassava germplasm in Africa. Results from other species demonstrate the lack of a clear grouping pattern of the germplasm based on phenotypic data alone [50, 51]. However, further studies should be conducted to establish a relationship between the clusters formed based on SNPs and morphological descriptors.

The use of SNP markers allowed us to identify which genotypes were definitely not duplicates, and identify putative duplicate accessions. To confirm true duplicates, we propose that high density genotyping, such as DArTSeq (Diversity Array Technologies) should be performed. The elimination of duplicate accessions should reduce the costs associated with conservation at the CGB in Côte d’Ivoire. We propose the adoption of the 36 SNP markers involved in this study for quality control at various stages of breeding process through varietal tracking using a unique fingerprint in cassava growing regions of Eastern and Western Africa.

## • Availability of data and materials

The datasets used and/or analyzed during the current study are available from the corresponding author on reasonable request.

## • Authors’ contributions

JSP, NKK and MEF initiated and designed the study. JSP and MEF mobilize the fund for the research. BN, NKK, DHO, EFY and WJLA collected samples. MEF developed the SNP markers. EFY, KMHK, NKK, TS, DHO and MEF analyzed data. MEF, EFY, WJLA, FS, DHO, and JMM wrote the manuscript. BN, MKK, LPLV-L, TS, RS, DK, SPAN and NY reviewed the manuscript. All authors read, corrected and approved the manuscript.

## • Acknowledgments

The authors thank the Centre National de Recherche Agronomique (CNRA) and the Centre Suisse de Recherche Scientifique (CSRS) for providing plant material.

## Supporting information

**S1 Fig. Genotype accumulation curve showing the minimum number of SNPs needed to differentiate all unique genotypes is seven.** The graph was developed using package Poppr in R software.

**S2 Fig. Population structure of the total set of the 95 cassava accessions assuming K = 2 (red and green) developed using STRUCTURE 2.3.4.** (A) Membership probabilities of each accession in order and (B) cluster membership probabilities of each accession sorted by Q (membership probabilities). Each accession is represented by a vertical bar

**S3 Fig. Inference of the number of K groups for the total set of the 95 cassava accessions according to Pritchard [35], as obtained using the program Structure Harvester [38].** The most probable number of genetic groups, two, is indicated by a red arrow. DeltaK = mean(|L”K)|)/sd(L(K)), L = Likelihood-log

**S4 Fig. Plot of Discriminant analysis of principal components (DAPC) showing the Bayesian Information Criterion (BIC) values indicating that the best number of clusters is five (red arrow) for the total set of the 95 cassava accessions**

**S5 Fig. Plot of Discriminant analysis of principal components (DAPC) for five assigned genetic clusters from the total set of the 95 cassava accessions, each indicated by different colors.** Dots represent different varieties. Inset left bottom corner and inset right bottom corner, show the eigenvalues of the 21 principal components and the eigenvalues of the first two discriminant function retained for the analysis respectively

**S6 Fig. Inference of the number of K groups for 77 retained cassava accessions according to Pritchard [35], as obtained using the program Structure Harvester [38].** The most probable number of genetic groups was three as indicated by a red arrow. DeltaK = mean(|L”K)|)/sd(L(K)), L = Likelihood-log.

**S7 Fig. Plot of Discriminant analysis of principal components (DAPC) showing the Bayesian Information Criterion (BIC) values indicating that the best number of clusters is three (red arrow) for the total set of the 77 retained cassava accessions**

**S8 Fig. Cluster membership probabilities of each accession based on the discriminant functions of the Discriminant analysis of principal components (DAPC) for the total set of the 77 cassava accessions.** Each accession is represented by a vertical bar. The membership coefficient of the accessions varied from 80% to 100%

**S9 Fig. Loading plots of Discriminant analysis of principal components (DAPC) showing the most contributing loci of the discriminant function (A) along axis 1 loci was Me.MEF.c.2574 (0.10) and (B) along axis 2 was Me.MEF.c.2268 (0.17)**

**S1 Table. The 95 cassava accessions from Côte d’Ivoire**

**S2 Table. The 34 cassava accessions from others regions of Africa**

**S3 Table. Genetic differentiation parameters by locus from the Côte d’Ivoire germplasm.** *Fst*, fixation index showing identity of individuals within sub-populations compared to those from other sub-populations within the total population; *Fis*, fixation index showing differentiation of individuals within sub-populations; *Fit*, fixation index showing homozygosity of individuals in the total population; **, *P* < 0.01; ***, *P* < 0.001

**S4 Table. Groupings of the 77 accessions following the Ascending hierarchical clustering, the ADMIXTURE program and the DAPC analysis**

## REFERENCES

[1] Hodgkin T, Bordoni P. Climate change and the conservation of plant genetic resources. J Crop Improv. 2012; 26: 329–345

[2] Mittal RK. ICAR–CGIAR. agric coop. 2017

[3] Maxted N, Kell S, Brehm JM., Jackson M, Ford-Lloyd B, Parry M. Crop wild relatives and climate change. Plant genetic resources and climate change. 2013; 291

[4] FAO, Produire plus avec moins: Le manioc guide pour une intensification durable de la production. 2013

[5] Ferguson ME, Shah T, Kulakow P, Ceballos H. A. global overview of cassava genetic diversity. PloS one. 2019;14: e0224763

[6] Glémin S, Scornavacca C, Dainat J, Burgarella C, Viader V, Ardisson M, et al. Pervasive hybridizations in the history of wheat relatives. Sci Adv. 2019; 5: eaav9188

[7] Mtunguja M, Ranjan A, Laswai H, Muzanila Y, Ndunguru J, Sinha N. Genetic diversity of farmer-preferred cassava landraces in Tanzania based on morphological descriptors and single nucleotide polymorphisms. Plant Genet Resour. 2017; 15: 138–146

[8] Benesi IRM,. Labuschagne MT, Hermeslan L, Mahungu N. Ethnobotany morphology and genotyping of cassava germplasm from Malawi. J Biol Sci. 2010; 10: 616–23

[9] Albuquerque HYGD, Oliveira EJD, Brito AC, Andrade LRBD, Carmo CDD, Morgante CV, et al. Identification of duplicates in cassava germplasm banks based on single-nucleotide polymorphisms (SNPs). Scientia Agricola. 2019; 76: 328–36

[10] Korir NK, Han J, Shangguan L, Wang C, Kayesh E, Zhang Y, et al. Plant variety and cultivar identification: advances and prospects. Crit Rev Biotech. 2013; 33: 111–25

[11] Elhoumaizi MA, Saaidi M, Oihabi A, Cilas C. Phenotypic diversity of date-palm cultivars (*Phoenix dactylifera* L.) from Morocco. Genet Resour Crop Evol. 2002; 49: 483–90

[12] Racchi ML, Bove A, Turchi A, Bashir G, Battaglia M, Camussi A. Genetic characterization of Libyan date palm resources by microsatellite markers. 3 Biotech. 2013; 4: 21–32

[13] N’Zué B, Okoma MP, Kouakou AM, Dibi KEB, Zohouri GP, Essis BS, et al. Morphological characterization of cassava (*Manihot esculenta* Crantz) accessions collected in the centre-west, south-west and west of Côte d’Ivoire. Greener J Agric Sci. 2014; 4: 220–31

[14] Djaha KE, Abo K, Bonny BS, Koné T, Amouakon WJL, Koné D, et al. Caractérisation agromorphologique de 44 accessions de manioc (*Manihot esculenta* Crantz) cultivées en Côte d’Ivoire. Int J Biol Chem Sci. 2017; 11: 174–84

[15] Kawuki RS, Herselman L, Labuschagne MT, Nzuki I, Ralimanana I, Bidiaka M, et al. Genetic diversity of cassava (*Manihot esculenta* Crantz) landraces and cultivars from southern, eastern and central Africa. Plant Genet Resour. 2013; 11: 170–81

[16] Yoon MS, Song QJ, Choi IY, Specht JE, Hyten DL, Cregan PB. BARCSoySNP23: a panel of 23 selected SNPs for soybean cultivar identification. Theor Appl Genet. 2007; 114: 885–99

[17] Schlötterer C. The evolution of molecular markers — just a matter of fashion. Nat Rev Genet. 2004; 5: 63

[18] Chagné D, Batley J, Edwards D, Forster JW. Single nucleotide polymorphisms genotyping in plants. Association mapping in plants. 2007; 77–94

[19] Oyesigye E, Zacarias A, Mondjana A, Magaia H, Ferguson M. Single nucleotide polymorphism (SNP) diversity of cassava genotypes in relation to cassava brown streak disease in Mozambique. Plant Genetic Resources; 2018; 16: 533–543

[20] Rabbi IY, Kulakow PA, Manu-Aduening JA, Danky AA, Asibuo JY, Parkes EY, et al. Tracking crop varieties using genotyping-by-sequencing markers: A case study using cassava (*Manihot esculenta* Crantz). BMC Genet. 2015; 16: 1–11

[21] Mtunguja M, Ranjan A, Laswai H, Muzanila Y, Ndunguru J, Sinha N. Genetic diversity of farmer-preferred cassava landraces in Tanzania based on morphological descriptors and single nucleotide polymorphisms. Plant Genetic Resources. 2017; 15: 138–146

[22] Ferguson ME, Hearne SJ, Close TJ, Wanamaker S, Moskal WA, Town CD, et al. Identification, validation and high-throughput genotyping of transcribed gene SNPs in cassava. Theor Appl Genet. 2012; 124: 685–95

[23] Das G, Rao GJN. Molecular marker assisted gene stacking for biotic and abiotic stress resistance genes in an elite rice cultivar. Front Plant Sci. 2015; 6: 698

[24] Hartl DL, Clark AG. Principles of population genetics. Sunderland, MA: Sinauer associates. 1997:116

[25] Thiruvenkadan AK, Jayakumar V, Kathiravan P, Saravanan R. Genetic architecture and bottleneck analyses of Salem Black goat breed based on microsatellite markers. Vet World. 2014; 7: 733–37

[26] Botstein D, White RL, Skolnick M, Davis RW. Construction of a genetic linkage map in man using restriction fragment length polymorphisms. Am J Hum Gen. 1980; 32(2): 314

[27] Wright S. Evolution and the genetic of populations. The theory of gene frequencies, Univ. Chicago Press. 1969; 2

[28] Wright S. Evolution and the genetic of populations. Variebility within and among natural populations. Univ. Chicago Press. 1978; 4: 295

[29] De Meeûs T, Goudet J. A step-by-step tutorial to use HierFstat to analyse populations hierarchically structured at multiple levels. Infect Genet Evol. 2007; 7: 731–35

[30] Nagy S, Poczai P, Cernák I, Gorji AM, Hegedus G, Taller J. PICcalc: An online program to calculate polymorphic information content for molecular genetic studies. Biochem Gene. 2012; 50: 670–72

[31] Jombart T, Devillard S, Balloux F. Discriminant analysis of principal components: A new method for the analysis of genetically structured populations. BMC Genet. 2010; 11: 94

[32] Kamvar ZN, Tabima JF, Everhart SE, Brooks JC, Krueger-Hadfield SA, Sotka E. Package ‘poppr’. 2019

[33] Dray S, Dufour A. The ade4 Package: Implementing the Duality Diagram for Ecologists. _Journal of Statistical Software. 2007; 22:1–20

[34] Larmarange J. JLutils: Collection of R functions. R package. 2021. Version 1.22.0. https://github.com/larmarange/JLutils

[35] Pritchard JK, Stephens M, Donnelly P. Inference of population structure using multilocus genotype data. Genet Soc Am. 2000; 155: 945–59

[36] François O, Durand E. Spatially explicit Bayesian clustering models in population genetics, Mol Ecol Resour. 2010; 10: 773–84

[37] Evanno G, Regnaut S, Goudet J. Detecting the number of clusters of individuals using the software STRUCTURE: a simulation study. Mol Ecol. 2005; 14: 2611–20

[38] Earl DA, Bridgett M. STRUCTURE HARVESTER: a website and program for visualizing STRUCTURE output and implementing the Evanno method. Conserv Genet Resour. 2012; 4: 359–61

[39] Jombart T, Collins C. Analysing genome-wide SNP data using adegenet 2.0. 0. 2015

[40] Oksanen J, Blanchet FG, Friendly M, Kindt R, Legendre P, McGlinn D, Minchin PR et al. Vegan: Community Ecology Package. 2020

[41] Wickham H. ggplot2: Elegant Graphics for Data Analysis. Springer-Verlag New York. 2016

[42] Sievert C. Interactive Web-Based Data Visualization with R, plotly, and shiny. Chapman and Hall/CRC Florida. 2020.

[43] Asare AP, Galyon IKA, Sarfo KK, Tetteh JP. Morphological and molecular based diversity studies of some cassava *(Manihot esculenta* Crantz) germplasm in Ghana. African J Biotechnol. 2011;13900–08

[44] Kizito EB. Genetic and Root Growth Studies in Cassava (Manihot esculenta Crantz): Implication fro Breeding. Acta Univ. Agric. Sueciae. 2006; 82: 1–127

[45] Elias M, McKey D, Panaud O, Anstett MC, Robert T. Traditional management of cassava morphological and genetic diversity by the Makushi Amerindians (Guyana, South America): perspectives for on-farm conservation of crop genetic resources. Euphytica. 2001; 120:143–57

[46] N’Zué B. Caractérisation morphologique, sélection variétale et amélioration du taux de multiplication végétative chez le manioc *(Manihot esculenta* Crantz (Euphorbiaceae). Thèse de doctorat. Université de Cocody, Département de Génétique. 2007

[47] Pissard A, Arbizu C, Ghislain M, Faux AM, Paulet S, Bertin P. Congruence between morphological and molecular markers inferred from the analysis of the intra-morphotype genetic diversity and the spatial structure of Oxalis tuberosa Mol Genetica. 2008; 132: 1–85

[48] Sawler J, Reisch B, Aradhya MK, Prins B, Zhong G-Y, Schwaninger H, et al. Genomics assisted ancestry deconvolution in grape. PLoS One. 2013;8(11), e80791

[49] Oliveira EJ, Ferreira CF, da Silva Santos V, de Jesus ON, Oliveira GAF, da Silva MS, Potential of SNP markers for the characterization of Brazilian cassava germplasm. Theor Appl Genet. 2014; 127: 1423–40

[50] Xia XC, Reif JC, Melchinger AE, Frisch M, Hoisington DA, Beck D, et al. Genetic diversity among CIMMYT maize inbred lines investigated with SSR markers: II. Subtropical, tropical midaltitude, and highland maize inbred lines and their relationships with elite U.S. and European maize. Crop Sci. 2005; 45: 2573–82

[51] Semagn K, Magorokosho C, Vivek BS, Makumbi D, Beyene Y, Mugo S, et al. Molecular characterization of diverse CIMMYT maize inbred lines from eastern and southern Africa using single nucleotide polymorphic markers. BMC Genom. 2012; 3: 113

